# Modeling cell turning by mechanics at the cell rear

**DOI:** 10.1101/2020.06.03.132456

**Authors:** Kun Chun Lee, Greg M. Allen, Erin L. Barnhart, Mark A. Tsuchida, Cyrus A. Wilson, Edgar Gutierrez, Alexander Groisman, Julie A. Theriot, Alex Mogilner

## Abstract

In this study, we explore a simulation of a mechanical model of the keratocyte lamellipodium as previously tested and calibrated for straight steady-state motility [1] and for the process of polarization and motility initiation [2]. In brief, this model uses the balance of three essential forces (myosin contraction, adhesive drag and actin network viscosity) to determine the cell’s mechanical behavior. Cell shape is set by the balance between the actin polymerization-driven protrusion at the cell boundary and myosin-driven retraction of the actin-myosin network. In the model, myosin acts to generate contractile stress applied to a viscous actin network with viscous resistance to actin flow created by adhesion to the substrate. Previous study [3] demonstrated that similar simple model with uniform constant adhesion predicts a rotating behavior of the cell; however, this behavior is idealized, and does not mimic observed features of the keratocyte’s turning behavior. Our goal is to explore what are the consequences of introducing mechanosensitive adhesions to the model.

## Myosin-powered retrograde actin network flow

Experimental and theoretical studies have established that myosin contracts actin arrays and generates contractile stress and that this stress grows with increasing myosin concentration [4, 5]. We make the simplest assumption that the myosin-generated contractile stress, *kM*, is linearly proportional to the myosin density, *M*. Here *k* is the proportionality coefficient (typical force per myosin unit) that in the model depends on blebbistatin/calyculin A treatment. The contractile force applied to the actin network is the divergence of the stress; in the case of the scalar stress, its gradient, *k*∇*M*. Following [6], we assume that adhesion complexes generate viscous resistance to the flow of F-actin (with velocity 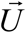 in the lab coordinate system). The respective resistance force, 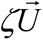, where *ζ* is the effective drag coefficient that we also refer to as adhesion strength, is balanced by the active contractile stress: 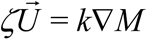.

The simple equation 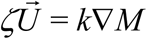 does not take into account passive stresses in the F-actin network due to its deformation during the flow. To add these passive stresses, we follow [6] and assume that these stresses have viscous character on the relevant time scale of tens of seconds. The small elastic component of the stress in the lamellipodium can be neglected [6], so we model a combination of the shear and deformation stresses in the F-actin with the formula 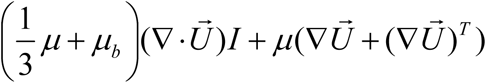, where *μ* and *μ*_*b*_ are the shear and bulk viscosities, respectively, and *I* is the identity tensor. Adding the divergence of these passive stresses to the myosin and adhesion forces results in the force balance equation determining the flow rate:

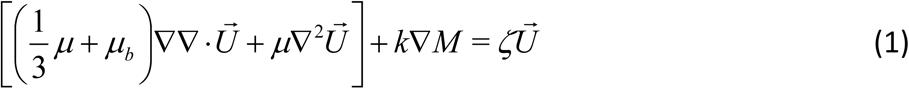

The boundary condition is the zero pressure at the free lamellipodial boundary:

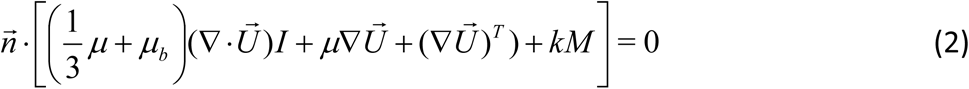

Here 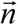 is the local normal unit vector to the lamellipodial boundary. The model assumes that the F-actin viscosity is spatially constant, independent of the F-actin density. Note, that due to this assumption we do not simulate and track actin density. Including a more detailed assumption of the viscosities being a function of the F-actin density does not change the qualitative pattern of the actin flow [6].

## Myosin transport

Following [6], we assume that myosin molecules bind and move with the F-actin network. Myosin molecules can detach from the F-actin, diffuse in the cytoplasm and reattach. Here, we assume that detachment and reattachment is rapid, in which case the system of equations for the actin-associated and diffusing myosin molecules [6] reduces to just one equation for the actin-associated myosin [1]. In this model, the rapid cycles of the detachment, diffusion in the cytoplasm and reattachment effectively result in a slow diffusion of the actin-associated myosin combined with the convective drift of myosin due to coupling with the F-actin that has a characteristic actin network velocity, 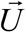 :

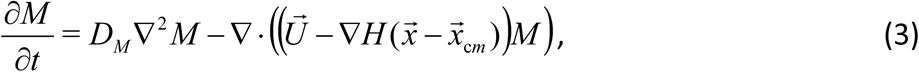

where *D*_*M*_ is an effective diffusion constant. The second term in Eq. (3) has an additional factor responsible for the observed expulsion of myosin from the center of the cell 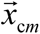 where thenucleus is located. This expulsion is achieved with introducing the smoothed Heaviside function:

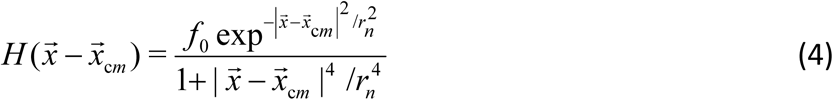

where *r*_*n*_ is effective radius of the nucleus and *f*_0_ is the effective repulsion strength. This function is approximately equal to *f*_0_ in the area covered by the nucleus and zero outside the nucleus. The gradient of this function introduces effective drift of myosin away from the nuclear center at the nuclear boundary.

The boundary conditions for the myosin transport are:

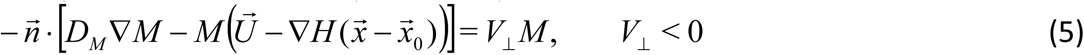

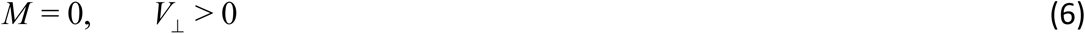

where 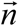 is the outward normal at the cell boundary. The left hand side of Eq. (5) is the total (diffusion-drift) flux of myosin at the retracting boundary. When the boundary is not moving (*V*_⊥_ = 0), Eq. (5) becomes the usual no flux boundary condition. When the boundary is moving inward (*V*_⊥_ < 0), additional inward myosin flux arises due to the fact that the total myosin is conserved, so the inward-moving cell edge collects the myosin (with local density *M*) at the edge and advects this myosin into the cell interior. To conserve the myosin density, this advection flux is the expression in the right hand side of Eq. (5). Eq. (6) describes the approximate no flux condition at the protruding boundary (*V*_⊥_ > 0). Due to effective diffusion, we have to use the total (diffusion-drift) flux of myosin at the protruding boundary. However, the effective diffusion is very slow. Thus, we can use the approximation that the myosin flux at the protruding boundary is equal to zero, which means *M* = 0 at this part of the boundary (Eq. 6). This approximate boundary condition, where we treat the minuscule myosin concentration at the protruding edge as zero, does produce a very small loss of conservation of total myosin density (typically < 0.02% of myosin per second is lost), so to restore the conservation of total myosin density we have added an additional step of uniformly re-normalizing the myosin density to each time step. This procedure amounts to assuming that there is a reservoir of unbound myosin that is in equilibrium with the bound pool.

## Mobile Cell Boundary

The cell boundary evolves according to the superposition of the inward boundary displacement from myosin-induced contraction and the outward displacement from actin polymerization. The net rate of boundary displacement,*V*_⊥_ in the locally normal direction is expressed in the model as:

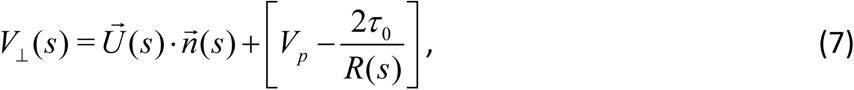

where *s* is the arclength parameter marking the position along the cell boundary, 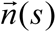 is the outward pointing locally normal vector and 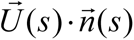 is the local actin centripetal flow given by Eq. (1) projected onto the normal to the boundary. The term in the square brackets is the net actin polymerization rate, where *V*_*p*_ is the polymerization rate, and the second term accounts for the effect of the membrane tension on decreasing the polymerization rate. Experimentally, it has been established that the lamellipodial area is conserved, likely due to a fixed amount of plasma membrane area [1]. As the membrane is effectively unstretchable, the membrane tension would increase as the cell area increases, decreasing the polymerization rate. We model this mechanics by assuming that *V*_*p*_ is constant along the boundary but decreases when the cell area increases. The second term in the square brackets is the effective Laplace pressure which prevents development of sharp corners at the boundary. This term is small and does not affect the global cell behavior, and scales with the local boundary curvature 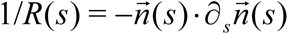. The proportionality coefficient 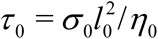 where *σ*_0_ is effective membrane tension (along the boundary), *l*_0_ is a length scale of molecular dimension and *η*_0_ is the effective drag coefficient for the membrane as the surface evolves. In essence, 2*σ*_0_ /*R*, is the Laplace pressure, multiplied by 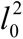 gives us the total force on a patch membrane of size 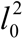. If we divide the total force 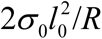 by the drag coefficient associated with the patch of membrane, then we get the velocity at which the patch of membrane is moving. Note that *τ* _0_ has the same dimension as a diffusion constant.

## Adhesion

Adhesion strength *ζ* appearing in Eq. (1) varies spatially [6,7] so 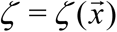. We model function 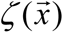 using patterns base on experimentally measured traction forces and distribution of adhesion molecules [7]. It is known that the traction force is weakest at the rear and strongest at the sides of cell. This implies that adhesion strength should be the greatest at the sides and the smallest at the rear and the strength of adhesion at the leading edge should be in between that of the rear and of the sides (Fig. 1A).

**Figure 1.**
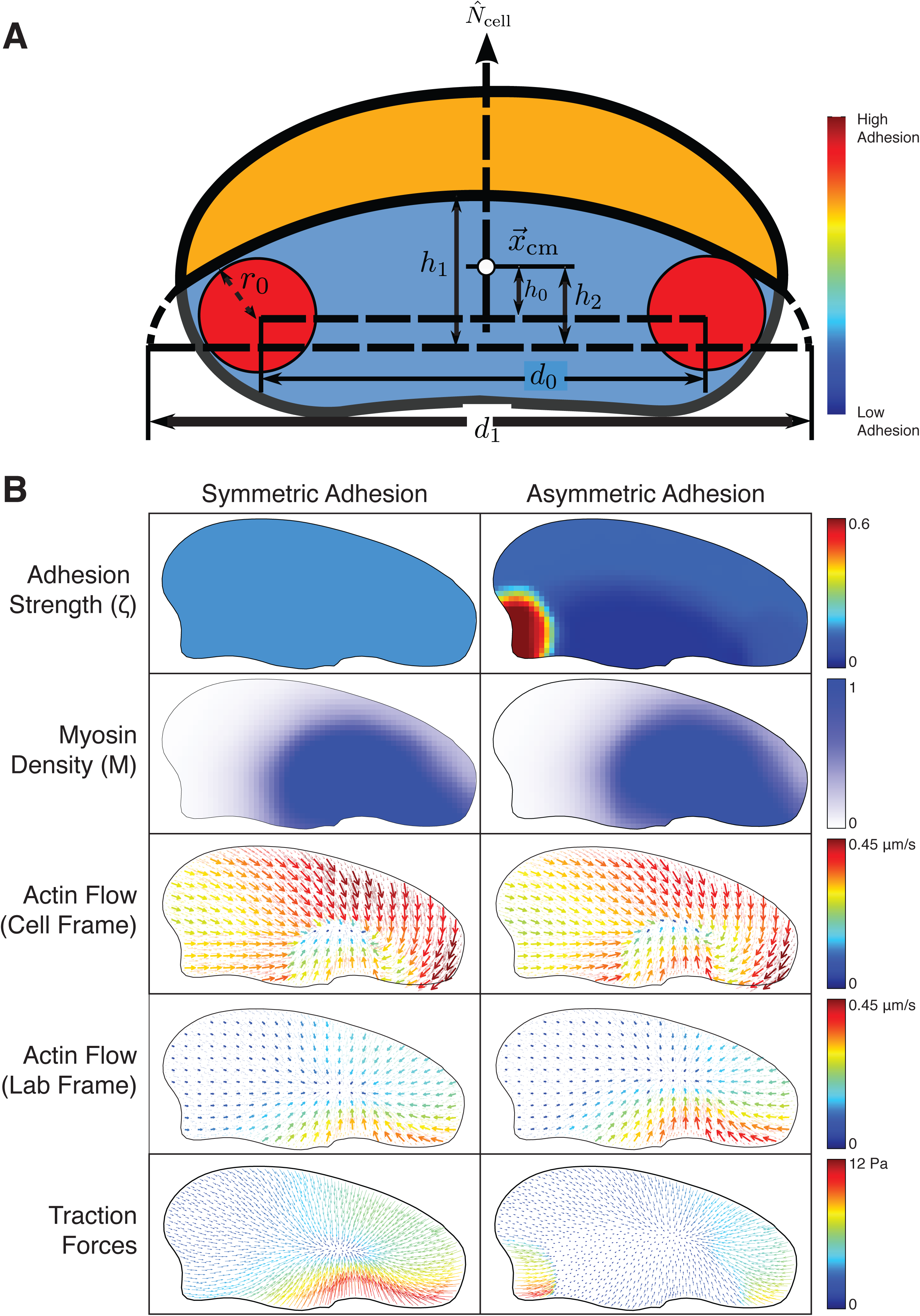
Simulations of a turning cell with fixed shape. (A) Schematic of the spatial distribution of adhesion strength. The two circles at the cell sides represent the regions where the adhesion strength is the highest (*red*). Adhesion strength is medium at the front (*yellow/orange*) and weakest at the rear (*blue*). (B) Detailed simulations of cytoskeletal asymmetries in the steadily turning cell with the fixed shape. In the simulations, adhesion distribution is pre-determined and is constant in time. Myosin distribution, actin flow and traction forces are computed according to the model dynamics. Left column: adhesion (*top*) is constant. Myosin (*2nd row*) is swept by the flow to the outer side of the cell. Both actin flow (*3*^*rd*^ *row*) and traction forces are high at the outer side of the cell. Right column: adhesion is moderate in the band along the leading edge, low at the cell rear, and high at the inner rear corner of the cell. Myosin is swept by the flow to the outer side and actin flow is high at the outer side of the cell with little change due to an asymmetric adhesion distribution.

To come up with a mathematical description for the pattern shown in Fig. 1A, it is best to work in the cell frame of reference and for convenience we can use the center of mass 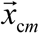 (center of the cell body in the model) as the origin of our coordinate system. To define the *x* and *y* axes, we first compute the eigenvectors of the gyration tensor as defined by *R*_*ij*_ = ∫*x*_*i*_ *y*_*j*_ d*A* where the integration is over the whole cell. Because this is a symmetric real tensor, it will diagonalize to create two unique orthogonal eigenvectors {*ê*_s_, *ê*_l_} as long as the cell is not circularly symmetric. These two eigenvectors by construction point along the longest and the shortest dimensions of the cell. We next construct a dynamical vector variable 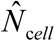 which acts like a compass that follows the shortest dimension (*ê*_s_) corresponding to the rear-front direction. This vector evolves according to:

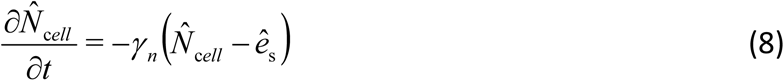

where 1/*γ* _*n*_ is the characteristic fast response time the exact value of which does not affect the predicted behavior. At the beginning of the run, we set 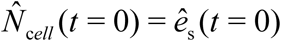 as our initial condition. Since 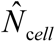 points along the short dimension of the cell we may designate it as the *y* axis and use the perpendicular line crossing 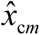 as the *x* axis. Using this coordinate system, we initially place two sites of locally maximal adhesion to the sides of the cell where the adhesion strength peaks are at the coordinates (−*h*_0_, ±*d*_0_ /2) (Fig. 1A).

To define the dynamic position of the adhesion peaks at the cell sides, we define a generalized Heaviside step function, keeping in mind that we are using the 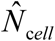 coordinate system:

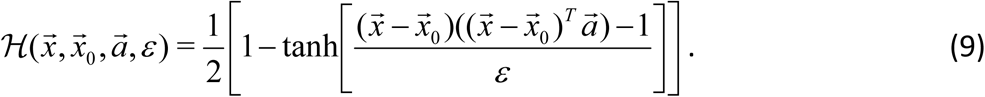

In this expression, vector quantities should be interpreted as column vectors so that 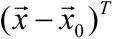 which stands for the transpose of 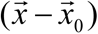 is a row vector. Just like the regular step function, the value of this function is 1 inside some region and 0 outside of this region with a transition zone width approximated by 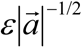. The shape of the region defined by this function is such that if the argument of the tanh function is negative/positive, then position 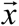 is inside/outside the given region, respectively. To see how this function works, notice that if we multiply out the vectorial quantities in the function argument using 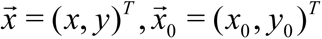 and 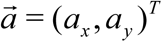 then

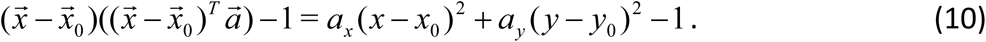

If we let *a*_*x*_ = *a*_*y*_ = 1/*r* ^2^, we get

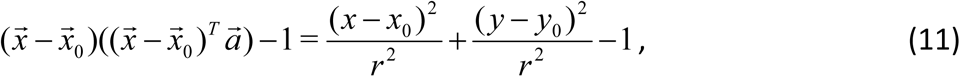

which tells us that the shape of the given region is a circle with radius *r* centered at 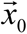. If *a*_*x*_ ≠ *a*_*y*_, then we have an ellipse.

We define *ζ* with the help of function ℋ and the coordinate system 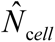 matches so that it closely with the adhesion pattern shown in Fig. 1A:

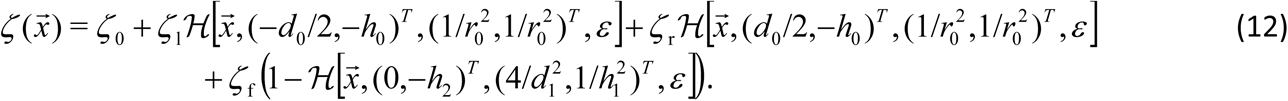

Here *ζ* _0_ is the baseline value for the adhesion strength (light blue region at the rear shown in Fig. 1A), *ζ* _f_ is the adhesion strength value at the leading edge, and *ζ* _l_ and *ζ* _r_ are the adhesion strength values at the left and right sides of the cell respectively.

### Varying adhesion strengths *ζ* _*l*_ and *ζ* _*r*_ in time

To test how adhesion asymmetry causes the cell to turn dynamically, we allow the adhesion strengths at the sides, *ζ* _l_ and *ζ* _r_, to oscillate in time according to equations:

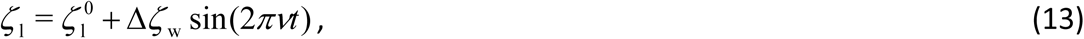

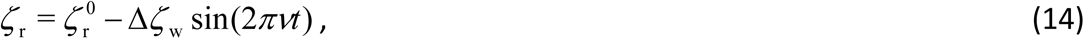

where 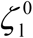 and 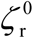 are the baseline adhesion strengths at the two sides. Δ*ζ*is the maximum deviation from *ζ* _w_and 1/*v* is the period of oscillation. We also used other functions of time that are always bounded from below and above) to model more stochastic variance of the adhesion strengths. For example, we used Ornstein-Uhlenbeck stochastic process (random walk in time of an overdamped harmonic oscillator perturbed by the white Gaussian noise). Results did not depend strongly on the nature of the time-dependent variation.

### Adhesion dynamics and boundary-crossing simulation

We modeled cells crossing boundaries of different adhesion strength, by first taking 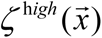 and 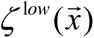 to denote the adhesion strength of the cell when the cell is crawling on high and low adhesivity substrate respectively. To smoothly let the cell cross between the two different substrates and thus transition between *ζ* ^h*igh*^ and *ζ* ^l*ow*^ (and reverse), we again use a different type of Heaviside step function ℒ which gives a value of one to the region in space with high adhesion and a value of zero to the areas with low adhesion substrate. Mathematically, this may be expressed as (using the lab coordinate system)

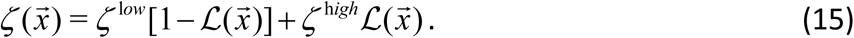

The precise form of ℒ is similar to that for ℋ (Eq. 9) with a different argument function.

### Simulation setup

The initial condition for our simulation was a circular cell of area *A* = 600*μ* m ^2^. We initially spread myosin uniformly over the whole cell. Symmetry was then broken by choosing a fixed orientation of the short axis of the cell, 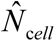 and biasing spatially the adhesion strength for the first minute minute, of the simulation. During this first minute, cells typically evolved into the characteristic crescent shape, after which 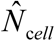 was allowed to evolve according to Eq(8).

All calculations are carried out using LGPL-licensed finite-element solver FreeFem++ (www.freefem.org) as described in detail in [1].

## Model parameters

The model variables and parameters are listed in the Tables below. Most of the parameter values are taken directly from our previous publications [1, 6] with minor changes. In the following sections, we discuss the physically relevant parameters.

### Viscous actin-myosin network

We take the characteristic length to be the typical cell size *L*_0_ = 10*μ* m [1] and the characteristic speed to be the characteristic cell speed, *V*_0_ = 0.2*μ* m/s [1] which is comparable to the retrograde flow rate of the actin network. The shear viscosity, *μ* = 5 kPa ×s [6], the bulk viscosity is normally higher than the shear viscosity, as the gels are more resistant to compression than shear so we use the value *μ*_*b*_ = 100 kPa ×s [6].

In order for the myosin stress to generate the observed flow of the order of *V*_0_ = 0.2*μ* m/s inside a lamellipodium with characteristic thickness of *h* ∼ 0.2*μ* m [6], the typical force scale is *μhV*_0_ = *f*_0_ = 200 *pN* : In our calculations, we multiply the viscosities by the characteristic thickness of the lamellipodium *h* = 0.2 m in order to convert the 3D stress derivatives into the 2D surface force densities. To non-dimensionalize Eq. (1), we choose 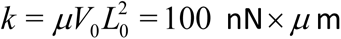 based on dimensional analysis. Using this coefficient *k*, 100 units of myosin in our scheme are expected to generate an average force density on the order of 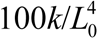 and hence capable of generating an average traction force on the order of 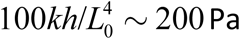, comparable to known traction force data [7, 8]. For all our simulation runs, unless stated otherwise, we use total amount of myosin *M* _t*otal*_ = 80 non-dimensional units. (Note that *M* _t*otal*_ is conserved in the model.) We set the diffusion coefficient *D*_*M*_ = 1.2 *μ* m ^2^ /s to be sufficiently small to keep the dimensionless Peclét number 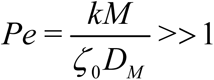 such that the actin flow dominates over diffusion [9] when the adhesion strength is minimal (*ζ* = *ζ* _0_).

### Myosin dynamics with fixed cell boundary

As part of our test calculation, we consider what happens when the shape of the cell is fixed (*V*_⊥_ = 0), but the cell is turning. To simulate such a situation, we solve Eq. (3) while taking the motion of the turning cell explicitly into account. This is done by adding a kinematic flow to Eq. (3) so that it becomes:

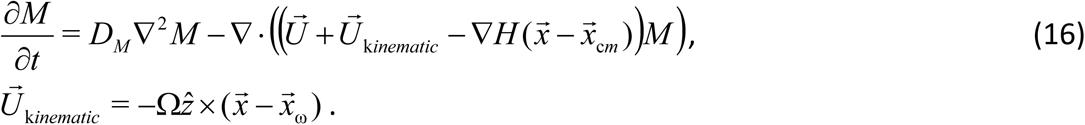

Here 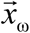 is the center of pivoting motion and 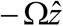 is the angular velocity of the cell. The negative sign accounts for the fact that myosin should drift in the direction opposite to the cell motion. 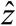 is the unit vector pointing out of the surface on which the cell crawls. For these computational runs, we choose 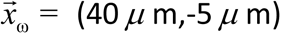 relative to the cell center-of-mass and Ω = 1/250 s ^−1^. These values mean that the cell is moving at a linear speed of 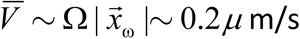 and angular speed about 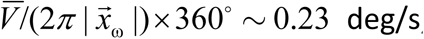, comparable to measured speed and angular speed in the experiments.

### Magnitude of adhesion strength

The strength of adhesion is characterized by the coefficients introduced in Eq. (12), namely {*ζ* _0_, *ζ* _l_, *ζ* _r_, *ζ* _f_}. We fix the baseline strength at *ζ* _0_ = 0.03 nN s/ *μ* m ^4^ for all of our runs. This value is comparable to the low adhesion strength we previously reported in [1]. The value for the other coefficients varies depending on what system we are studying but we try to keep them all comparable to the ‘medium’ values that we have reported previously 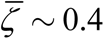. Note that with the ‘medium’ values of adhesion strength as the characteristic retrograde flow rate in our simulation of 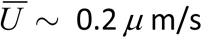, the characteristic traction force is 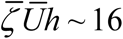 Pa which is comparable to the experimentally measured traction force (5-10 Pa).

### The nucleus

In our two-dimensional model the nucleus is represented as a disc with radius *r*_*n*_ = 7.5*μ* m centered at the cell center-of-mass 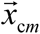. To effectively repel both myosin and adhesion from the region where the nucleus resides, we need to choose 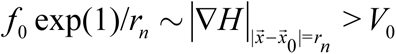 for the repulsion to be strong enough to counteract the actin flow. We choose *f* = 1*μ* m ^2^ /s, which yields *f*_0_ exp(1)/*r*_*n*_ ∼ 0.35*μ* m/s.

### Cell shape dynamics

The dynamics of the cell boundary is dictated by the balance of net local protrusion/retraction rate and the Laplace pressure. The strength of the Laplace pressure term is governed by 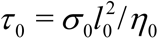. We choose tension *σ*_0_ to be ∼ 0.1nN/ *μ* m [10]. The drag coefficient for the membrane *η*_0_ scales, according to Stoke’s Law, as ∼ 6*πμ*_0_*l*_0_, where *μ*_0_ = 1cP is the viscosity of water. As for *l*_0_, it should be the size of lipid molecule, approximately 1nm. Using these numbers, *τ* ∼ 0.5*μ* m ^2^ /s. In this study, we used a comparable value of *τ* = 0.1*μ* m ^2^ /s, which means that the tension is slightly smaller than 0.1nN/ *μ* m.

### Dependence of the model behavior on the parameters

The most important parameters in the model are the myosin strength *k*, the adhesion strength *ζ*, the actin viscosity *μ*, and the characteristic cell speed*V*_0_. The model behavior is sensitive to these parameters in the sense that for the shape and movement of the model cell to resemble the real cell, there is a number of constraints on these parameters that have to be in place, analyzed in detail in [1, 6]. These constraints are not rigid: a fewfold changes of these parameter values (Table 3) still predicts qualitatively all the observed behavior. Similarly, the model is robust to a few fold change of all characteristic adhesion strengths used (Table 4). The model is even less sensitive to all other parameters listed in Table 3: changes of up to an order of magnitude of their values (one at a time, of course) do not change the predicted behavior qualitatively. We emphasize that most of the parameters orders of magnitude are known from extensive studies of keratocyte cells.

**Table 1:** Model variables.

| Variable | Meaning | Dimension |
| --- | --- | --- |
| $t$ | time | s |
| $s$ | arc length | $\mu\text{ m}$ |
| $\vec{x}$ | two-dimensional coordinate | $\mu\text{ m}$ |
| $M(\vec{x}, t)$ | myosin concentration | units/$\mu\text{ m}^3$ |
| $\vec{U}(\vec{x})$ | local F-actin flow velocity | $\mu\text{ m/s}$ |
| $\vec{n}(\vec{x})$ | local normal unit vector to the lamellipodial edge | non-dimensional |
| $\zeta(\vec{x}, t)$ | adhesion strength/drag-coefficient | nN s/$\mu\text{ m}^4$ |
| $V_{\perp}(s)$ | net local protrusion/retraction rate | $\mu\text{ m/s}$ |
| $V_p$ | local polymerization rate | $\mu\text{ m/s}$ |
| $R(s)$ | local radius of curvature | $\mu\text{ m}$ |

**Table 2:** Definition of adhesion strength parameters

| Parameter | Meaning | Dimension |
| --- | --- | --- |
| $\zeta_0$ | baseline adhesion strength | nN s/$\mu\text{ m}^4$ |
| $\zeta_l$ | adhesion strength coefficient for the left wing | nN s/$\mu\text{ m}^4$ |
| $\zeta_r$ | adhesion strength coefficient for the right wing | nN s/$\mu\text{ m}^4$ |
| $\zeta_f$ | adhesion strength coefficient for the leading edge (front) | nN s/$\mu\text{ m}^4$ |
| $\zeta_l^0$ | baseline adhesion strength for the left wing | nN s/$\mu\text{ m}^4$ |
| $\zeta_r^0$ | baseline adhesion strength for the right wing | nN s/$\mu\text{ m}^4$ |
| $\Delta\zeta$ | maximum amplitude of adhesion strength perturbation | nN s/$\mu\text{ m}^4$ |
| $\nu$ | frequency of modulating adhesion strength | $\text{s}^{-1}$ |
| $h_0$ | y-displacement relative to the cell center for the side adhesion patches | $\mu\text{ m}$ |
| $h_1$ | front-back length parameter for front adhesion patch size | $\mu\text{ m}$ |
| $d_0$ | x-displacement relative to the cell center for the side adhesion patches | $\mu\text{ m}$ |
| $h_1$ | width parameter for front adhesion patch size | $\mu\text{ m}$ |
| $r_0$ | side adhesion patch size | $\mu\text{ m}$ |
| $\varepsilon$ | adhesion transition zone width parameter | dimensionless |

**Table 3:**
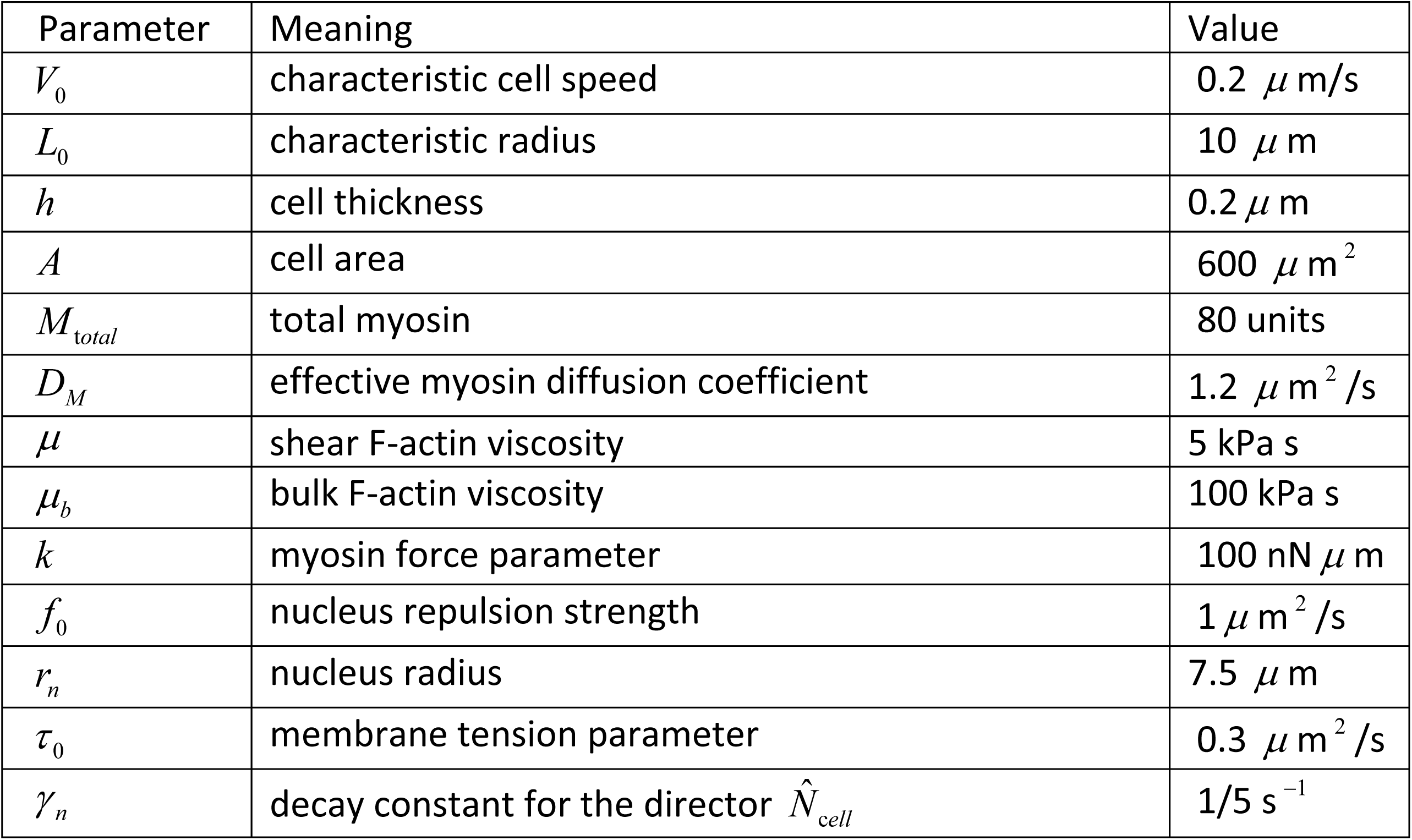
Non-adhesion related constant model parameters.

| Parameter | Meaning | Value |
| --- | --- | --- |
| $V_0$ | characteristic cell speed | $0.2 \mu\text{m/s}$ |
| $L_0$ | characteristic radius | $10 \mu\text{m}$ |
| $h$ | cell thickness | $0.2 \mu\text{m}$ |
| $A$ | cell area | $600 \mu\text{m}^2$ |
| $M_{total}$ | total myosin | 80 units |
| $D_M$ | effective myosin diffusion coefficient | $1.2 \mu\text{m}^2/\text{s}$ |
| $\mu$ | shear F-actin viscosity | 5 kPa s |
| $\mu_b$ | bulk F-actin viscosity | 100 kPa s |
| $k$ | myosin force parameter | $100 \text{nN} \mu\text{m}$ |
| $f_0$ | nucleus repulsion strength | $1 \mu\text{m}^2/\text{s}$ |
| $r_n$ | nucleus radius | $7.5 \mu\text{m}$ |
| $\tau_0$ | membrane tension parameter | $0.3 \mu\text{m}^2/\text{s}$ |
| $\gamma_n$ | decay constant for the director $\hat{N}_{cell}$ | $1/5 \text{s}^{-1}$ |

**Table 4:** The values of adhesion strength parameters used

| Parameter | Standard | High Adhesion | Low Adhesion | Mild Turn | Tight Turn |
| --- | --- | --- | --- | --- | --- |
| $\zeta_0 [\text{nNs}/\mu\text{m}^4]$ | 0.03 | 0.03 | 0.03 | 0.03 | 0.03 |
| $\zeta_f [\text{nNs}/\mu\text{m}^4]$ | 0.4 | 0.4 | 0.12 | 0.4 | 0.4 |
| $\zeta_l [\text{nN s}/\mu\text{m}^4]$ | 0.5 | 0.5 | 0.15 | N/A | N/A |
| $\zeta_r [\text{nNs}/\mu\text{m}^4]$ | 0.5 | 0.5 | 0.15 | N/A | N/A |
| $\zeta_l^0 [\text{nNs}/\mu\text{m}^4]$ | N/A | N/A | N/A | 0.5 | 0.5 |
| $\zeta_r^0 [\text{nNs}/\mu\text{m}^4]$ | N/A | N/A | N/A | 0.5 | 0.5 |
| $\Delta\zeta [\text{nNs}/\mu\text{m}^4]$ | N/A | N/A | N/A | 0.1 | 0.45 |
| $\nu [\text{s}^{-1}]$ | N/A | N/A | N/A | 0.5 | 0.5 |
| $h_0 [\mu\text{m}]$ | 2 | 2 | 2 | 2 | 2 |
| $h_1 [\mu\text{m}]$ | 13 | 13 | 13 | 13 | 13 |
| $d_0 [\mu\text{m}]$ | 14 | 14 | 14 | 14 | 14 |
| $d_1 [\mu\text{m}]$ | 22 | 22 | 22 | 22 | 22 |
| $r_0 [\mu\text{m}]$ | 2.5 | 2.5 | 2.5 | 2.5 | 2.5 |
| $\varepsilon [\text{dim/less}]$ | 0.25 | 0.25 | 0.25 | 0.25 | 0.25 |

## Simulation results

### Asymmetries in myosin distribution, actin flow and traction forces in turning cell

We first considered a cell with a fixed shape (Fig. 1B), chosen to approximate an experimentally measured turning cell shape (Fig. 3f in [11] and [12]). Similar to the observed turning cells, there is a lower aspect ratio on the slower side and a higher aspect ratio on the faster side, i.e. the outer wing is more elongated than the inner wing. For simplicity, in these simulations we removed the nucleus and its effects from the cell center. With the fixed cell boundary, we used the kinematic actin-myosin flow of a turning cell as explained above in Eq. (16), as well as a fixed adhesion distribution in time. We tested two distributions of adhesions to see which could replicate observed measurements of actin flow, myosin distribution and traction forces. In one simulation, adhesion was constant in space, while in the other it varied in space with adhesion higher on the inner side of the turning cell. The adhesion distribution with resulting simulation results are presented in Fig. 1B. We observed that the myosin distribution in both cases was biased toward the fast side of the cell due to the kinematic actin flow, with only slight insignificant differences in myosin distributions between these two cases. However, only an asymmetric adhesion distribution could replicate experimental measurements of traction forces [12].

**Figure 2.**
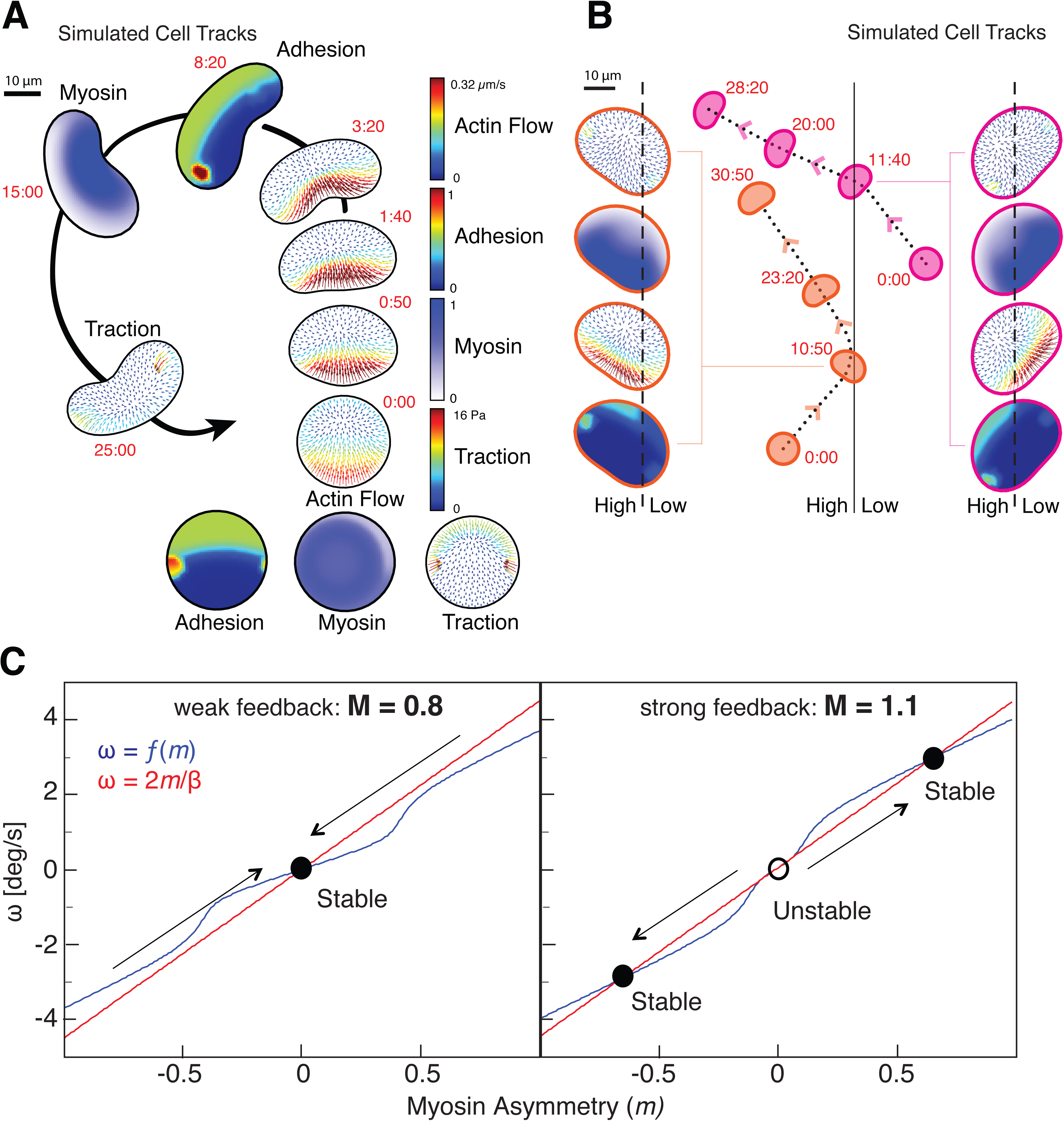
Characteristics of cell turning can be reproduced with a mechanical model. (A) The results of a free-boundary simulation of cell migration where cell shape and migration evolved with the initial assumption of increased adhesion strength on the left side of the cell. Cell outlines are shown at time points as labeled in minutes (*red*), with the first two time points not presented in their actual location in space. Note that we observe expected changes in cell shape, actin flow (*3:20*), adhesion asymmetry (*8:20*), myosin distribution (*15:00*) and traction forces (*25:00*). (B) Simulation with free-boundary model of cell migration of cells migrating from high to low adhesion (*orange contours*) and from low to high adhesion (*magenta contours*). Cell shape and parameters at the point of crossing are presented on the sides as labeled, with scale bars as in panel A. Matching experimental observations [12], cells turn towards the side of the substrate with higher adhesion in both cases. Scale bar indicates 10 *µ*m. (C) From the combination of the linear relationship between angular speed and myosin asymmetry dictated by asymmetric delivery of myosin II to the outer side of the cell inducing turning (*red*) and the non-linear relationship between angular speed and myosin II asymmetry dictated by the weakening of adhesion by myosin II contractility according to a stick-slip model (*blue)*, the stable turning state of a cell can be determined. When myosin contractility is too weak to break adhesion (M = 0.8) then a single stable solution can be obtained as would be seen for a meandering cell (*left)*. When myosin contractility is stronger (M = 1.1) and weakens adhesions promoting further turning, then a cell will be stable in either of two persistent turning states (*right)*.

**Figure 3.**
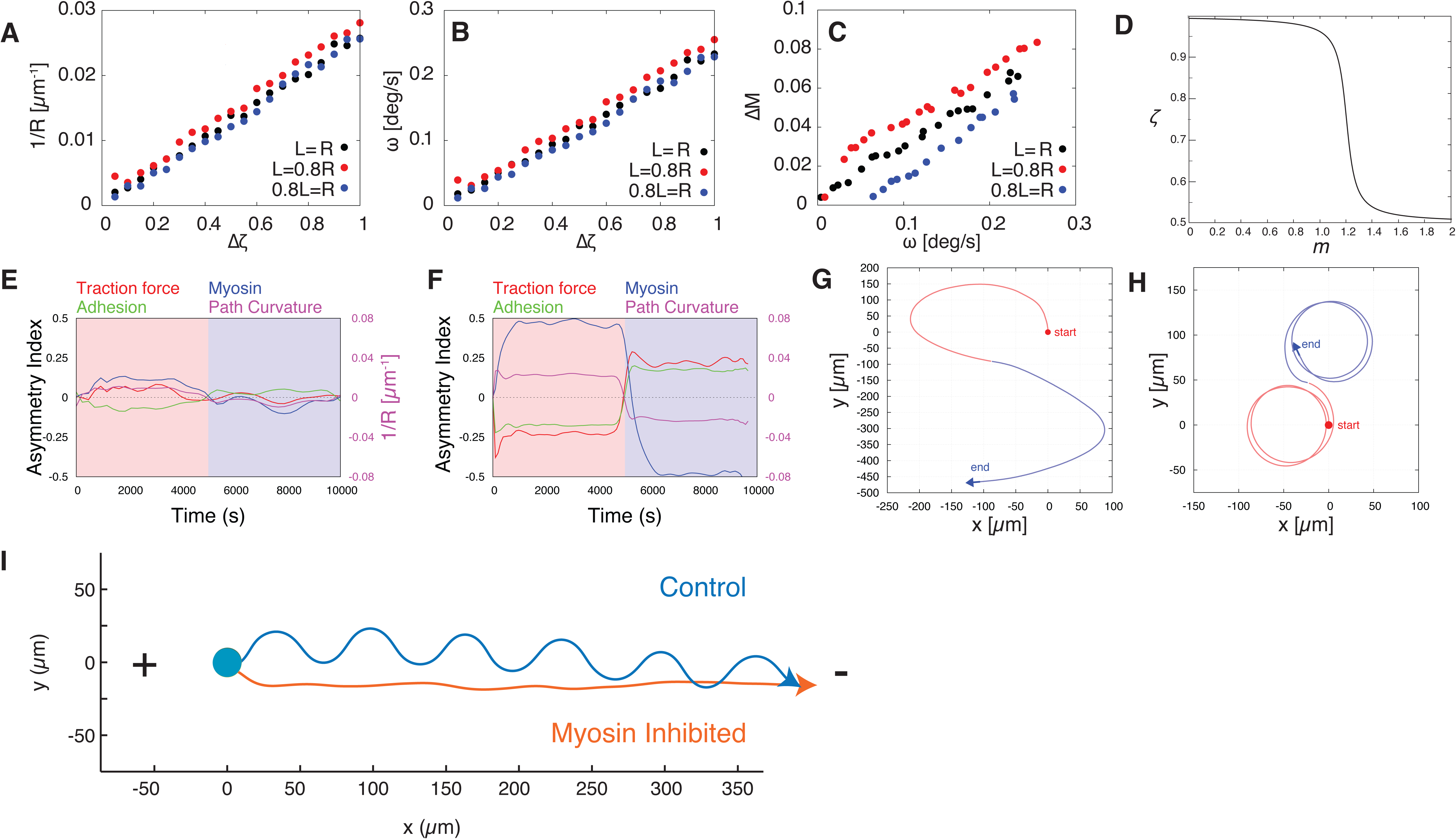
Modeling of asymmetries in adhesion and incorporation into a comprehensive model reproducing cell turning behavior. (A-C) For the free boundary model of cell migration the side-to-side difference in adhesion strength (Δ*ζ*) was fixed and in panel E the radius of curvature of the path (*R*), in panel F the angular velocity of the cell (*ω*), and in panel G the side-to-side difference in myosin concentration were calculated as the cell turned to the left. For each simulation adhesion strength at the front was either kept constant from side to side (black points), set as the left being 80% of the right (*red points*), or set as the right being 80% of the left (*blue points*). Differences of myosin concentration and adhesion strength are measured in units of average myosin concentration and adhesion strength, respectively. Note that changing adhesion strength at the leading edge made little contribution to cell turning. (D) A plot of the assumed adhesion strength, *ζ*, as a function of myosin concentration, *M*, where 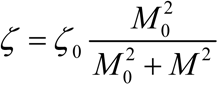. Increasing myosin concentration and hence contractility is assumed to decrease the effective adhesion strength according to a non-linear stick-slip model of adhesion. (E-H) The results of a free-boundary simulation of cell migration where cell shape and migration evolved with an initial assumption of increased adhesion strength on the left side of the cell (*pink zone*) and after 5,000 seconds with reversed adhesion asymmetry (*blue zone*). The normalized left-right asymmetry of traction forces, adhesion strength, myosin density as well as the path curvature are plotted as a time series for a single simulation in (*E*) with myosin contractility having no effect on adhesion strength, and (*F*) with increased myosin contractility negatively regulating adhesion strength. Tracks for each of these simulations are presented in (*G*) and (*H*) respectively. Each cell starts at the position marked with a red circle and migrates toward the blue arrow. Without the negative feedback of myosin contractility on adhesion strength, turning is unstable and lower in magnitude. With this negative feedback included in the simulation turning is higher in magnitude and more persistent. The normalized asymmetry is defined as(⟨*X*_*L*_⟩ − ⟨*X*_*R*_⟩) / ⟨*X*⟩. (I) Simulated trajectories of cells in an electric field under control conditions (*blue*) and with inhibition of myosin contractility (*orange)*.

### The model reproduces characteristic cell turning behavior

We used the free-boundary model of the cell to simulate the evolution of the cell from the symmetric stationary disc-like shape when initially the adhesion at the left is increased (Fig. 2A). First, we used the model in which the adhesion relaxes to the uniform, constant adhesion. In this case, the symmetric shape of the cell moving along the straight trajectory evolved (data not shown). Then, we simulated the model with the mechanosensitive adhesion, as described above. The steady asymmetric cell shape and turning with a constant-radius trajectory evolved from this initial condition, with the shape and patterns of myosin distribution, actin flow and traction forces (Fig. 2A) mimicking those observed in the experiment [12].

### Boundary crossing and myosin asymmetry

We used the free-boundary model of the cell to simulate the cell movement while crossing a boundary between high and low adhesions, as described above. In the simulations at each time step, we scaled the adhesion strength pattern depicted in Fig. 1A by a factor dependent on the parts of the cell that were on higher or lower adhesion, respectively. The results are shown in Fig. 2B. We found that, matching experimental results, cells would turn towards the side of higher adhesion after an asymmetry in adhesion developed at the cell rear. We also simulated the situation where the density of myosin was increased on one side of the cell (data not shown). The simulation demonstrated that an asymmetry in myosin produces a steady turn away from the side of increased myosin. Yet, once the externally imposed bias in myosin concentration is removed, myosin re-distributes around the cell, the cytoskeletal symmetry is restored, and the cell starts to move straight. This matched experimental findings from asymmetric exposure to the myosin activating small molecule calyculin (Figure 6A in [12]). The results indicate that the positive feedback between the kinematics of turning and myosin distribution are sufficient for transient but not persistent turning.

### Turning behavior with myosin-adhesion feedback

As the computations with the free-boundary model were computationally expensive, we developed the following combined analytic-computational theory to examine the long time scale cell trajectories. We first fixed the difference Δ*ζ* between the left and right adhesion strengths. As a result, the cell, after a brief transient relaxation, started to move with steady shape and angular speed, *ω*, along the trajectory with constant radius of curvature, *R*. Also as a result, a steady difference in myosin concentrations Δ*M*, between the left and right parts of the cell developed. We repeated such simulations varying the value of Δ*ζ* from 0 to the maximum (no adhesion at one side) and measured Δ*M, ω* and *R* for each adhesion asymmetry (Fig. 3A-C). We found a strong relationship between the asymmetry in adhesion and the predicted rate of turning and myosin asymmetry. In comparison, when we varied the adhesion strength at the leading edge, we found that such variance did not affect cell turning behavior significantly (Fig. 3A,B).

We found that the angular speed is an approximately linear function of the left-right adhesion difference: *ω* ≈ *α*Δ*ζ*, and the myosin concentration difference in the left and right sides of the cell is an approximately linear function of the angular speed: Δ*M* ≈ *βω* (Fig. 3A-C). Here*α* and *β* are defines as constant parameters identified in the simulations.

We then followed the experimental and theoretical findings in [2] of the negative feedback between local contraction and adhesion strength. In [2], the adhesion strength was the function of the local actin flow rate, but as the flow rate *V* is proportional to the ratio of the myosin concentration to the adhesion strength, here for simplicity we assume that the adhesion strength is the following function of the myosin concentration: 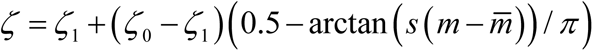, so that the adhesion strength is a decreasing function of the myosin concentration *m* (Fig. 3D). This is also consistent with the observed and predicted biphasic relationship between actin retrograde flow speed and traction stress [13], and the hypothetical “molecular-clutch” model of adhesions [14]. At the inner rear of the turning keratocyte, inward actin flow is slow and the molecular clutch of adhesions is in place, creating large traction forces. At the outer rear of the turning keratocyte, inward actin flow is high secondary to myosin contractility and the molecular clutch of adhesions fails, leading to small traction forces. To appropriately model our data, it is important that the adhesion strength decreases slowly at low myosin densities, faster at moderate myosin densities and slower again at high myosin densities, as in Fig. 3D. However, the exact functional form of this dependence is not critical, and we also note that such functional dependencies are ubiquitous in biology [15]. In the *ζ* (*m*) relation,*ζ* _0_ is the high adhesion strength (Table 4), and *ζ*_1_ = *ζ* _0_ /2 is the low adhesion strength, *s* = 20 is the parameter determining how fast the adhesion drops at threshold myosin density 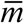 [2]; we choose 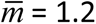 where *M* = 1 corresponds to average myosin density at the cell rear in the state of the straight movement.

According to this assumption of negative feedback of myosin contractility on adhesion strength, if the cell moves with angular speed*ω*, then there will be the resulting difference in myosin concentrations at the cell left and right sides, *ω*, then due to the resulting difference in myosin concentrations at the cell sides, *M* + *m* and *M* − *m*; 2*m* ≈ *βω*. Then, there will be the following side-to-side difference in the centripetal flow: *V*_*l*_ ∼ (*M* + *m*) / *ζ* _*l*_, *V*_*r*_ ∼ (*M* − *m*) / *ζ* _*r*_, where 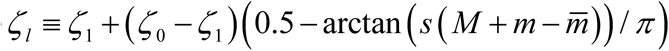 and 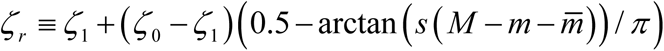. The resulting angular velocity is proportional to the difference (*V*_*l*_ −*V*_*r*_), and so:

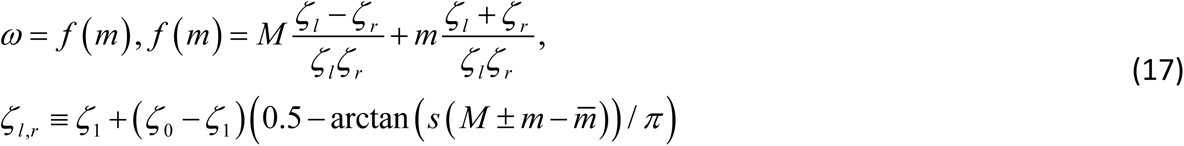

On the other hand, we have:

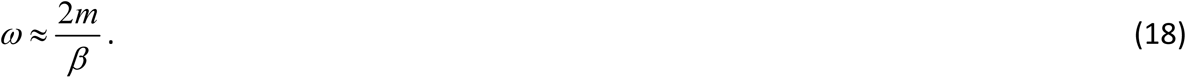

The system of Eq. (17,18) is shown graphically in Fig. 7C and the intersections of the two relationships determines the steady turning state of a cell. We investigate the behavior of this model when the average myosin density (or strength) (parameter *M*) changes. In this figure, we can see that for greater myosin strength, there are three steady states, one of which corresponds to straight migration and equal adhesion at the sides (*ω* = 0, Δ*ζ* = 0), and two others correspond to finite angular speeds and adhesion strength differences at the sides. These two finite angular speeds are the same in magnitude and opposite in sign, and they correspond to rotation in the clockwise and counter-clockwise directions. The movement with *ω* = 0 is unstable, while the two other ones are stable, so a cell with this described negative feedback between local myosin concentration and adhesion strength will switch between turning persistently in two opposite directions, matching the behavior of experimentally observed trajectories [12]. On the other hand, when myosin strength is low, the only stable state corresponds to straight migration and equal adhesion at the sides (*ω* = 0, Δ*ζ* = 0).

In order to illustrate that point, we numerically solved the system of two dynamic equations:

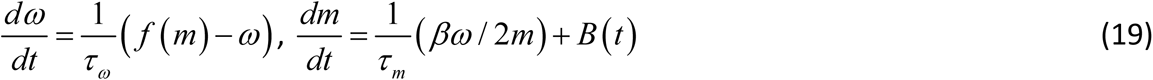

for the angular speed and adhesion strength difference that produce the steady solutions given by Eq. (17) and (18), and describe the relaxation of the angular speed and myosin difference to their steady states with characteristic times *τ*_*ω*_ and*τ* _*m*_, respectively. In the second equation of Eq. (19), the term *B* (*t*) is used to describe stochastic uncorrelated noise. Numerically, we can solve Eq. (19) using the Forward Euler method as follows:

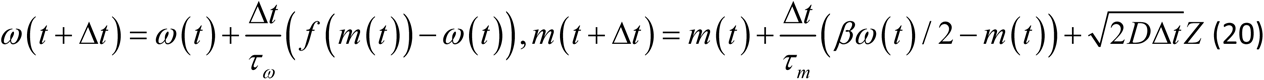

Here in the stochastic term *D* is the effective diffusion (random steps of adhesion change) of adhesion, and *Z* is the standard normal random variable. Note, that the same results can be achieved if the angular speed is noisy, or both speed and adhesion are noisy, with the magnitude of the noise adjusted to fit the results.

We simulated Eq. 20 numerically using parameters 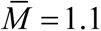 for control value of the myosin strength and 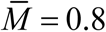 for low value of the myosin strength, *D* = 0.01, *β* = 0.45 (other parameter are listed above) and recorded the time series of cell angular speed (Fig. 2C). When the parameter *M* > 1, the negative myosin-adhesion feedback is strong enough to provide the bistable cell switching behavior (Fig. 2C) that results in peaks in the angular speed distribution, which correspond to persistent turning with rates on the order of ∼ 1 degree per second, as observed [12]. Simulations produce trajectories illustrated in Figure 3, which agrees with experimental results [12].

Finally, the experiment with cell motion in the electric field [12] showed that the cell in general moved along the direction to the cathode, but the directionality oscillated, so the cell followed a sinusoidal trajectory (Fig. 3I). The model reproduces this observation as follows. We assume that the electric field tends to orient the leading edge of the cell in the direction of the cathode. We model this tendency by adding the term −*rθ* to the equation for the rate of change of the angular velocity, where*θ* is the angle at which the cell moves relative to the cathode direction. Essentially, the angular velocity of the cell rear is slowed down by the leading edge pull proportionally to the deviation from the cathode direction when the cell is turning away from this direction, and accelerated when the cell is approaching this direction. Respective system of equation is:

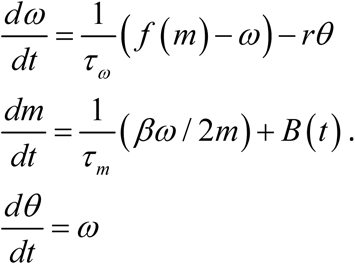

Numerical solutions of these equations with *r* = 0.3 and all other parameters the same as above are shown in (Fig. 3I) for both weak (*M* = 0.8) and strong (*M* = 1) myosin contractility.

## Summary

Free-boundary mechanical model of the keratocyte lamellipodium, with mechanosensitive adhesion and positive feedback between cell movement, myosin distribution and actin network flow, reproduces experimentally observed cell turning behavior.

